# The Divergence of Mean Phenotypes Under Persistent Directional Selection

**DOI:** 10.1101/2022.09.27.509694

**Authors:** Archana Devi, Gil Speyer, Michael Lynch

## Abstract

Numerous organismal traits, particularly at the cellular level, are likely to be under persistent directional selection across phylogenetic lineages. For such traits, unless all mutations affecting such traits have large enough effects to be efficiently selected in all species, gradients in mean phenotypes are expected to arise as a consequence of differences in the power of random genetic drift, which varies by approximately five orders of magnitude across the Tree of Life. Prior theoretical work examining the conditions under which such gradients can arise focused on the simple situation in which all genomic sites affecting the trait have identical and constant mutational effects. Here, we extend this theory to incorporate the more biologically realistic situation in which mutational effects on a trait differ among nucleotide sites. Pursuit of such modifications lead to the development of semi-analytic expressions for the ways in which selective interference arises via linkage effects in single-effect models, which then extend to more complex scenarios. The theory developed clarifies the conditions under which mutations of different selective effects mutually interfere with each others’ fixation, and shows how variance in effects among sites can substantially modify and extend the expected scaling relationships between mean phenotypes and effective population sizes.

---

Much of evolutionary biology relies on comparisons of mean phenotypes from distantly related species, followed by downstream attempts to develop plausible hypotheses for the observed patterns, almost always in the context of adaptive explanations. As phylogenetic lineages become isolated, their mean phenotypes are expected to diverge as a consequence of varying selection pressures. However, under many circumstances substantial divergence can be expected even in the face of identical selection pressures, owing to the vagaries of mutation and random genetic drift. In particular, by altering the accessibility of mutations to selection, a change in effective population size (*N*_*e*_) modifies the fixation probabilities of alternative alleles, with small *N*_*e*_ reducing the accumulation of beneficial alleles and increasing that of detrimental alleles. This leads to the expectation that there can be gradients in the performance of traits across species experiencing identical selection pressures, provided that the effects of all mutations are not so large as to be equally visible to natural selection at all population sizes (Lynch 2018, 2020).

Here we explore the consequences of a key determinant of the drift barrier to the mean performance of traits that has been ignored in prior theory development – the effects of a distribution of sites with varying effects on the phenotype. A substantial fraction of earlier work on the evolution of mean phenotypes assumes an infinite-alleles and/or infinite-sites model, whereby each newly arising mutation arrives at a site previously fixed in the population, while also assuming an absence of limits to the potential range of phenotypic variation (Kimura and Crow 1964; Kimura 1969; Latter 1970; Lande 1975; Bulmer 1980; Lynch and Hill 1986). Owing to its relative mathematical tractability, this model has played a central role in many areas of population genetics, including the development of theory on the maintenance of variation, the long-term response to selection, and the accumulation of deleterious mutations in various contexts (reviewed in Walsh and Lynch 2018).

However, for a wide variety of problems, the infinite-sites model is unrealistic biologically, and its utility as an approximation remains unclear. The concerns are numerous. First, the mutational target sizes of the molecular/cellular constituents of phenotypic traits are quite constrained in size. For example, an average protein is of order 1 kb in length, and specific functional domains generally encompass < 20 amino acids. Many elements at the level of DNA (e.g., transcription-factor binding sites) and RNA (e.g., microRNAs, and stems and loops of larger RNAs) are substantially smaller. The sizes of effectively nonrecombining linkage groups are often in the range of a few bp to several kb depending on the recombination rate. Second, the mutation rate is sufficiently high that in large populations, multiple independent mutations will often cosegregate at individual nucleotide sites, which can confer no more than four allelic types. Finally, the infinite-sites model has the undesirable property that the mutation spectrum is independent of the genetic background, resulting in a situation in which mean phenotypes can diverge without limits by either drift or directional selection. In reality, as more nucleotide sites in a stretch of DNA are occupied by deleterious mutations, the segment-wide deleterious and beneficial mutation rates must, respectively, decline and increase.

The approach taken here assumes a finite number of genomic sites contributing to the expression of a trait, with mutations at different sites potentially having different magnitudes of phenotypic/fitness effects, e.g., amino-acid replacement sites with different functional consequences for the encoded protein, silent sites under varying levels of selection owing to effects on mRNA folding and/or translational speed or accuracy, and noncoding sites with varying effects on gene expression. There has been growing interest in this type of model (Cockerham 1984; Charlesworth and Jain 2014; John and Jain 2015; Lynch 2018, 2020), but many problems remain to be solved.

Linked sites with differing mutational effects can be expected to play a significant role in phenotypic divergence owing to the multiple ways in which they interfere with each other in the selective process. For example, beneficial mutations at sites with small effects will be unavailable to selection if they arise in tight linkage with a segregating deleterious mutation at a site with large effects (Nguyen Ba et al. 2019). On the other hand, if sites with small effects greatly exceed the number of major-effect loci, beneficial mutations at the latter positions will have reduced visibility to selection if they happen to arise on a relatively poor linked background associated with segregating minor-effect sites. More generally, one can expect moderate-effect sites to experience both types of problems, particularly if there is an inverse relationship between the numbers of sites and their contributing effects. The overall process is further complicated by the fact that recurrent purging of deleterious mutations has general effects on effective population sizes, thereby influencing all other aspects of the efficiency of selection. There has been much research on these matters as well (Gerrish and Lenski 1998; Johnson and Barton 2002; Campos and Wahl 2010; Desai and Fisher 2007; Charlesworth 2013a; Good et al. 2014; Pénisson et al. 2017; Jain 2019), but almost all analyses have been restricted to the infinite-sites model, and often to populations that are effectively infinite in size with all mutations having equal effects.

## The Model

We start with a simple model with *L* linked sites (factors), each with two alternative allelic states, + and −, contributing positively and negatively to the trait, but with the magnitude of +*/*− effects allowed to vary among sites (Figure 1). Such a model would apply, for example, to a situation in which there is one optimal nucleotide at a site, with the remaining three having equivalent fitness effects. Because the stretch of nucleotide sites under consideration is assumed to be completely linked, the positions of the sites are irrelevant, and there can be a multiplicity of functionally equivalent haplotypes (i.e., with identical numbers of + alleles) in each effect class, which alters their ease of mutational accessibility (Lynch 2018, 2020). The site-specific per-generation mutation rates from the − to the + states, and vice versa, denoted as *u*_01_ and *u*_10_, respectively, will be assumed to be identical at all sites.

**Figure 1.**
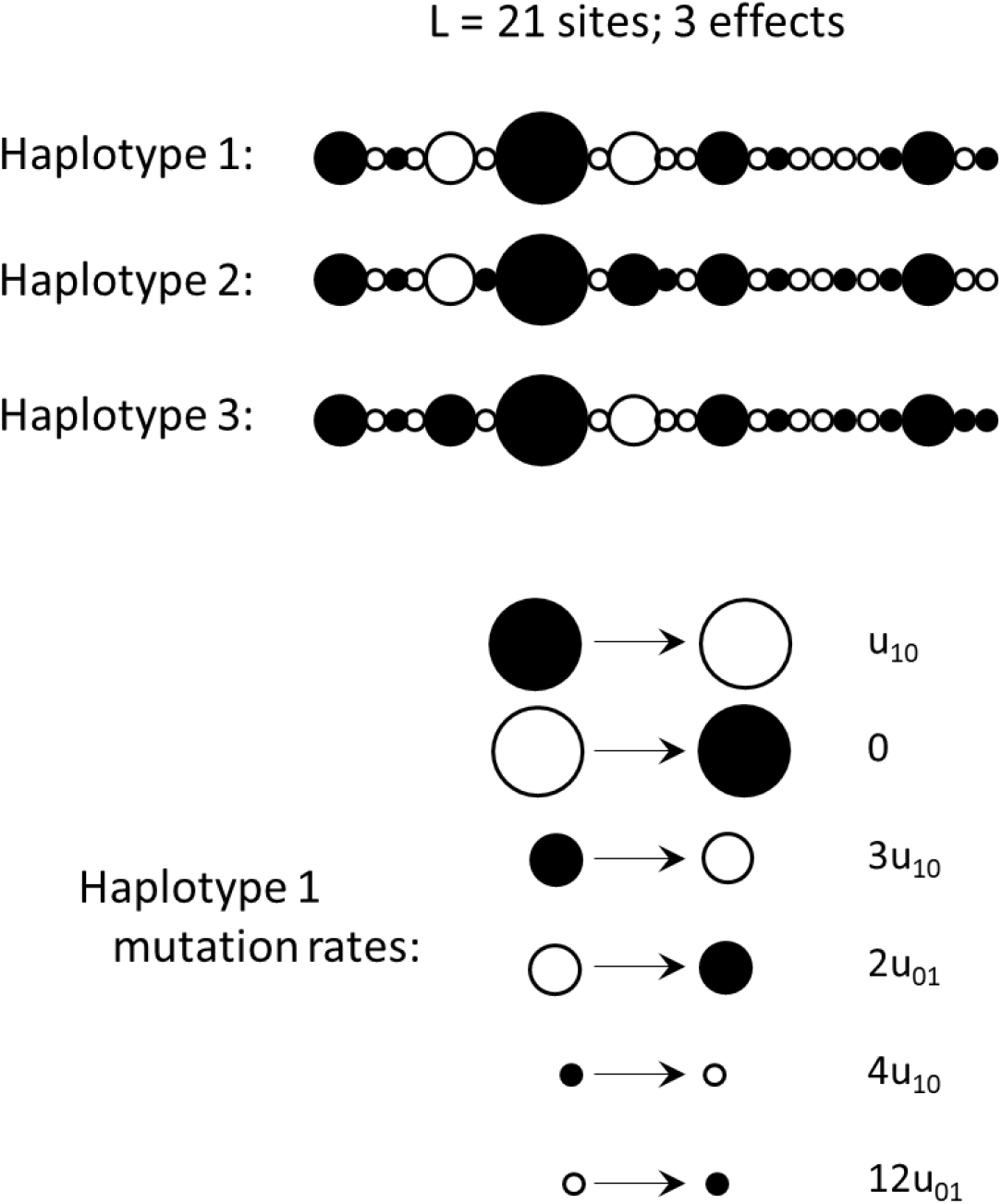
Schematic for the general approach. Here, there is a linkage block (experiencing no recombination) containing 22 sites, with an approximately exponential distribution of numbers of sites with three effects (one site with major effects, surrounded by five of medium effects, and 16 of small effects). Three of the many possible haplotypes are shown, with solid and open balls denoting + and − alleles. Given the assumption of complete linkage, the ordering of site-specific haplotypes is irrelevant, and in this case, haplotypes 2 and 3 are functionally equivalent, as they contain identical numbers of sites with + alleles for the three types of effects. The pattern of mutation is haplotype dependent, being a function of the numbers of + and − alleles at each type of site; the rates of the total set of possible mutations for haplotype 1 a.re given in the bottom panel, with *u*_01_ and *u*_10_ being the mutation rates from − to + allelic states, and vice versa. It is assumed that the site-specific mutation rates are low enough that individuals incur no more than a single mutation per generation.

As a central goal is to determine the conditions under which gradients in mean phenotypes can be expected under persistent directional selection in populations of different sizes, it is desirable to perform analyses with biologically realistic combinations of parameter values. Across the Tree of Life, *N*_*e*_ generally falls in the range of 10^4^ to 10^9^, and the mutation rate per nucleotide site scales negatively with the ∼ 0.76 power of *N*_*e*_ (Lynch et al. 2016; Long et al. 2017; Walsh and Lynch 2018). Thus, where computational work was involved, the following analyses were performed under the assumption of a deleterious-mutation rate per site (which might be a cluster of adjacent nucleotides) of 10^*−*7^ at an adult population size of *N* = 10^4^, such that *u*_10_ = 0.00011*N*^*−*0.76^, which is approximately 10× the known rate per nucleotide site. With this scaling, for the full range of population sizes employed here (*N* = 10^4^ to 10^9^), the product *Nu*_10_ then ranges from ≃ 0.01 mutations/population/site/generation at the lowest to 0.10 at the highest population sizes. It should be noted that the negative scaling of the mutation rate with absolute population size (*N*) is likely shallower than that assumed here, as *N*_*e*_*/N* for large multicellular species (small *N*_*e*_) is likely on the order of 0.1, whereas that for microbial species can be orders of magnitude smaller. In the end, we provide analytical approximations that make no assumptions about the relationship between mutation rates and population sizes.

We evaluate the consequences of a wide range of linkage-block lengths, from 1 (free recombination) to 10^6^, selection coefficients from *s* = 10^−8^ to 10^−4^, and mutation biases towards beneficial alleles *β* = *u*_01_*/u*_10_ = 0.10 to 1.00. Under this finite-sites model, the deleterious mutation rate per haplotype increases linearly with the number of sites harboring advantageous alleles, whereas the beneficial rate scales in the opposite direction.

The absolute population size consists of *N* haploid individuals, so that *de novo* mutations have initial frequencies of 1*/N*. Assuming independent fitness effects within and between loci (i.e., no dominance or epistasis), as done below, all results should extend to diploids by substituting 2*N* for *N* and 2*N*_*e*_ for *N*_*e*_. As noted below, the effective population size (*N*_*e*_), which is ≤ *N*, governs the magnitude of random genetic drift, and is a natural outcome of the structure of the linkage group, the strength of selection, and *N* itself.

The following work is performed under the assumptions of a classical Wright-Fisher discrete-generation model with sequential episodes of mutation, selection, and random genetic drift. Under this model, allele frequencies fluctuate in time, but because mutations are reversible, the system always eventually evolves to a quasi-steady-state distribution, provided the fitness function remains constant. Our particular focus is on how long-term average frequencies of beneficial alleles at various site types depend on the number and distribution of site types within linkage groups, on the joint forces of selection and mutation bias, and in particular on the population size. Related analyses have been performed by John and Jain (2015), Jain and John (2016), and Jain (2019), but mostly under the assumptions of either an effectively infinite population and/or an infinite-sites framework, and even in these cases, achieving reasonably simple expressions has been difficult.

Owing to the stochastic nature of the underlying processes, computer simulations of these processes must proceed for very large numbers of generations to achieve stable estimates of means and variances. To obtain greater computational speed, for large population sizes, we scaled the input parameters so as to keep *Nu*_10_, *Nu*_01_, and *Ns* constant, by reducing *N* and increasing the mutation and selection parameters by the same factor, with constraints such that *N* was always ≥ 10^3^, and *s* and *Lu*_10_ always ≤ 0.1. Burn-in periods before compiling statistics were typically at least 10^5^*N* generations, with the populations then being assayed every *N/*10 generations for 10^6^ to 10^8^ intervals. Simulations, which often extended for several days, were carried out with a program written in C++ (freely available from the authors), in a form that allows parallel analysis of multiple population sizes. Although we have evaluated a broad range of population-genetic environments extensively by computer simulation, throughout we attempt to provide heuristic semi-analytical expressions to address more general issues.

## Results

### Sites with single effects

For baseline comparisons, we start with simplest situation of sites with single effects, such that all beneficial mutations compete maximally with each other. Expanding on prior work (Lynch 2020), new expressions are presented to explain the general consequences of this extreme setting. The fitness function is assumed to be of the form 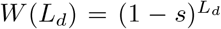, where *L*_*d*_ is the number of deleterious mutations carried in a haplotype, such that a maximum fitness of 1.0 occurs in individuals free of deleterious alleles, whereas with *L* equivalent sites, (1 − *s*)^*L*^ is the minimum fitness (for a haplotype containing only deleterious alleles). Under this multiplicative fitness model, selection operates on each site independently, and there is no epistasis. Although this leads to the expectation of no linkage disequilibrium in populations that are infinite in size (Eshel and Feldman 1970), this is not the case in finite populations.

The case of linkage blocks of length *L* = 1 is of special interest, as it represents the limiting situation of free recombination, where selection is most efficient. For this situation, an analytical expression for the long-term mean frequency of the + allele, here denoted 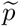, has already been developed by Kimura et al. (1963), and will not be repeated here, except to say that the fit to simulated data is excellent across the full range of population sizes, selection coefficients, and mutation rates. Although highly accurate, two undesirable features of the Kimura et al. (1963) solution are the need to solve a confluent hypergeometric function by a series expansion and the rather nontransparent interpretation of the formulations, and various approximations for particular domains of *Nu*_10_ and *Ns* have been given by Charlesworth and Jain (2014).

An alternative expression, which is quite accurate over the full range of parameter space explored herein and extends to larger linkage blocks, can be obtained in the following way. In Lynch (2020; Equation S10), it was noticed that if the within-population variance in numbers of mutant alleles per individual is known from simulations, the long-term average frequency of + alleles is accurately described by

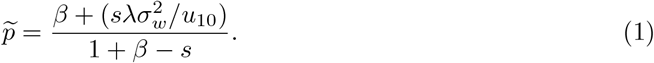

Here, 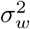 is the mean within-population variance per locus (i.e., the total variance in number of + alleles per individual divided by *L*), and *λ* = 1 − (1*/N*_*e*_) is a measure of the resistance of the population to random genetic drift, with *N*_*e*_ being the effective population size (Lynch et al. 1993). Derived from a quantitative-genetic perspective, this expression evaluates 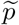 as the mean allele frequency at which the selection advance per generation (a function of the genetic variance) is matched by the decline associated with mutation. Others have used such a matching approach to estimate the position of the leading edge of the full distribution (Goyal et al. 2012; John and Jain 2015).

Letting 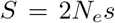, by extension from McVean and Charlesworth (1999) and Long et al. (2019),

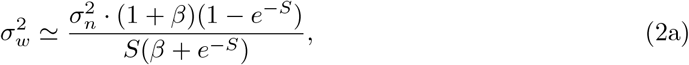

where

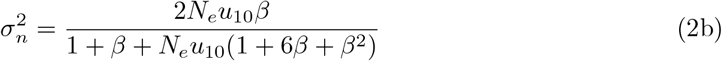

(from Lynch 2020; Equation S9) is the expected variance under neutrality (equivalent to half the expected neutral heterozygosity per site). A key remaining issue is that unless 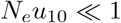, the effective population size (*N*_*e*_), will be depressed below the absolute population size (*N*), by selective interference among simultaneously segregating mutations. As a consequence, Equation 1 cannot be solved by substituting *N*_*e*_ = *N*, and a separate expression is needed for *N*_*e*_.

There are many ways to define an effective population size, depending on the allelic behavior of interest. One common consideration is the variance effective population size, i.e., the degree to which nucleotide diversity is depressed at neutral sites linked to other sites under selection (e.g., Charlesworth et al. 1995; Kim and Stephan 2000; Good et al. 2014; Campos and Charlesworth 2019). However, application of estimates of *N*_*e*_ obtained from simulations of standing levels of variation at linked neutral sites to the preceding formulae yields a less than satisfactory fit to observed levels of variation and mean allele frequencies.

An alternative approach starts with a consideration of the expected mean frequency of beneficial alleles over sites under the assumption of no interference (Li 1987; Bulmer 1991), and given the selection and mutation pressure (*s* and *β*), estimates the *N*_*e*_ necessary to account for an observed equilibrium beneficial-allele frequency, 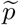 (Lynch 2020). The ratio *N*_*e*_*/N* is then

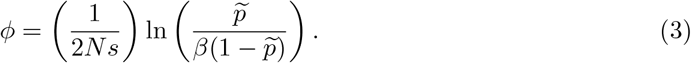

Using estimates of 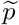 from simulated data to solve for *ϕ*, and substituting *N*_*e*_ = *ϕN* in the preceding expressions, Equations 2a,b provide excellent fits to observed within-population variances for the full range of parameters explored here, usually well within 10% of observed values (Supplemental Figure 1), whereas Equation 1 yields estimates of 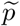 that are always within 3% of observed values (Figure 2). Notably, this approach was found to be valid for all linkage-block lengths explored, from *L* = 1 to 10^6^. Note also that as *Ns* → ∞,

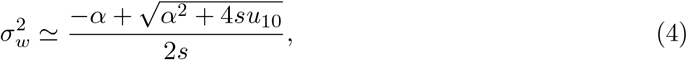

where *α* = *s* + *u*_01_ + *u*_10_. In this case, provided *s* exceeds the site-specific mutation rates (*u*_01_ and *u*_10_), 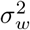 also closely approximates the equilibrium frequency of a deleterious allele in an infinite haploid (fully recombining) population, and more generally 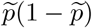.

**Figure 2.**
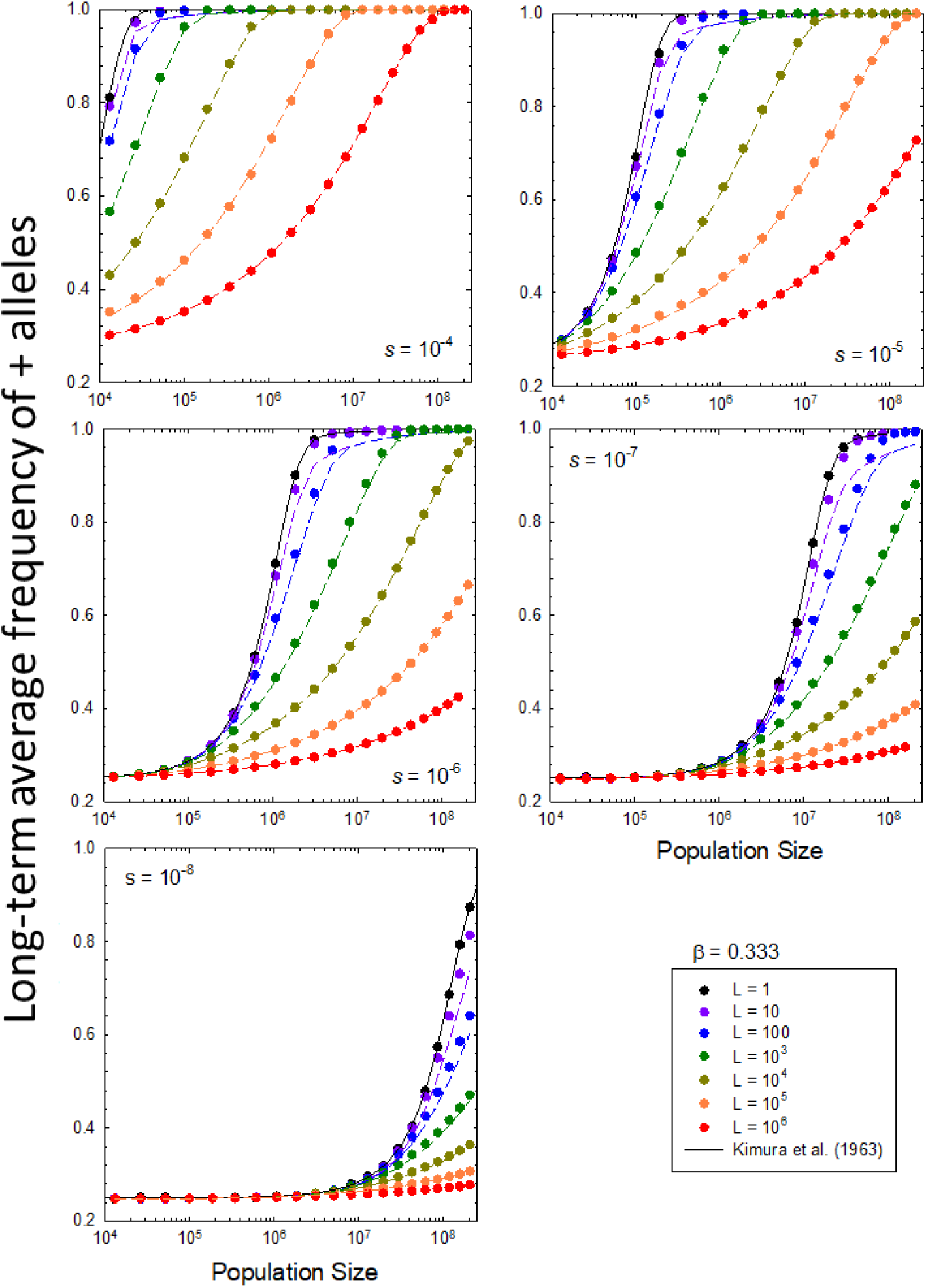
Relationship between the average frequency of advantageous alleles and the population size *N* (x axis), selection coefficients (different panels), and linkage-block length *L* (colored lines within panels) under the assumption of equal fitness effects across loci. The lines associated with each set of points are the theoretical predictions obtained with Equations 1, 2a, and 2b, using the fixation *N*_*e*_ derived from computer simulations. The mutation bias in all panels is *β*= 0.33, so the neutral expectation at small *N* is 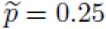.

Although these observations justify the use of the correction factor *ϕ* to transform *N* into the fixation effective population size relevant to equilibrium allele frequencies, validation of this approach required the use of estimates of *ϕ* derived by computer simulations. For more practical applications, we require an expression for *ϕ* from first principles. An excellent approximation to *ϕ*, as a function of the mutation rates, selection coefficient, number of loci, and absolute population size, was obtained by inspection in Lynch (2020), albeit with a particular scaling between the mutation rate and population size. In the Supplemental Text, we derive more general expressions, accounting for the amount of selective interference imposed on the fixation probability for beneficial mutations by linked sites.

Despite the complexity of the underlying issues, the derived expressions for *ϕ* generally yield estimates that are within 30% (often considerably closer) of simulation results (Figure 3). Although this is not a fully satisfactory outcome, given that *ϕ* varies 10,000-fold over the full range of parameter space, the essence of the system is captured. This provides an upgrade to the visual-fit interpretation of Lynch (2020), yielding insight into the scaling relationships between *ϕ* and the underlying population-genetic parameters. For example, for *L >* 10^4^,

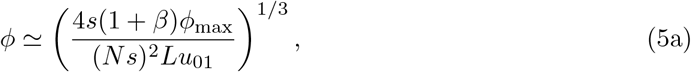

with

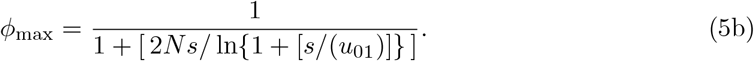

**Figure 3.**
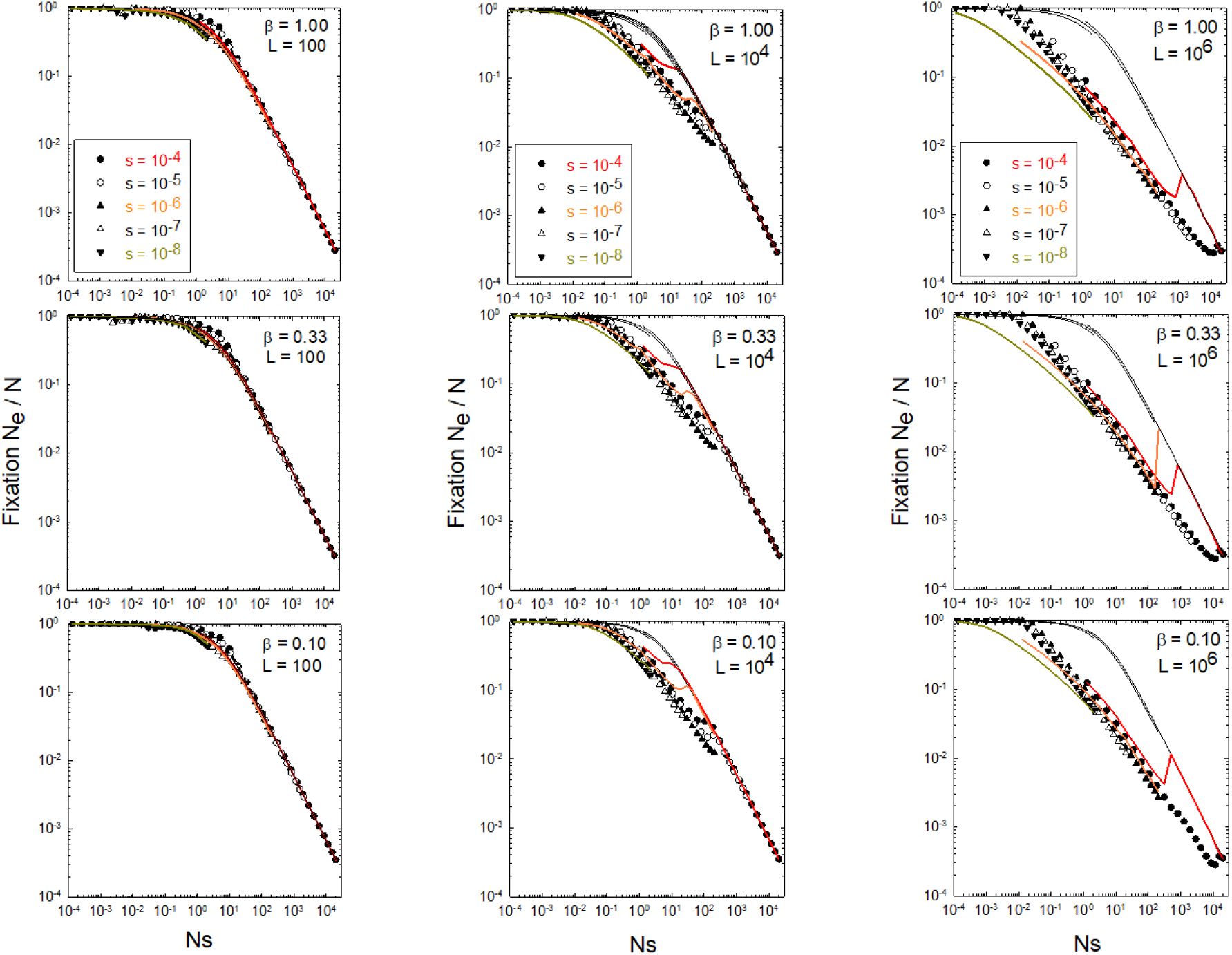
*ϕ* = *N*_*e*_*/N* as a function of *Ns* for three different linkage-block lengths *(L* = 10^2^, 104, and 10^6^; columns) and three levels of mutation bias *(β* = 1.0, 0.33, and 0.1; rows). The data points were obtained after applying estimates of 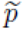 from computer simulations to Equation 3, whereas the colored lines (for three values of *s* in each panel) were obtained using expressions derived in the Supplemental Text; here solid lines are the full solutions to a transcendental equation, whereas the dashed lines are obtained from first-order approximations (closed expressions) – Equation All for *L* = 100 and Equation A10 for *L* = 10^4^ and 10^6^.

This expression shows that the reduction in *N*_*e*_ caused by linkage scales inversely with the cube root of the number of sites. It also shows that *ϕ* is a function of two other key composite parameters: the ratio of the selection strength to the mutation rate to beneficial alleles, and the ratio of the selection strength to the power of drift in the absence of interference, i.e., for large *L*,, *ϕ* scales with the ∼ 1*/*3 power of *s/u*_01_, and with the −2*/*3 to −1 power of *Ns* with increasing *Ns*.

Summing up for the simplest situation in which all sites within a linkage block have equivalent effects on fitness, contrary to the single-site expectations (Kimura et al. 1963), where there is a quantum shift in the frequency of beneficial alleles with increasing population size around a pivot point of *Ns* = 1, linkage reduces the gradient of response of 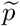 to *N* (Figure 2). Instead of a shift from the neutral expectation of 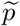 to that expected under deterministic selection-mutation balance over a window of just an order of magnitude of *N*, linkage can extend the gradient to several orders of magnitude of *N*, with the effect becoming increasingly pronounced with larger *L*. On the other hand, when viewed as a function of *N*_*e*_, where the latter is derived from the heterozygosity segregating at linked neutral sites, 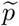 is largely (but not entirely) a stepwise function of 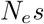 as *N*_*e*_ subsumes the influence of linkage interference. There is, however, some additional influence of *L* in the region of 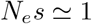 (Figure 4).

**Figure 4.**
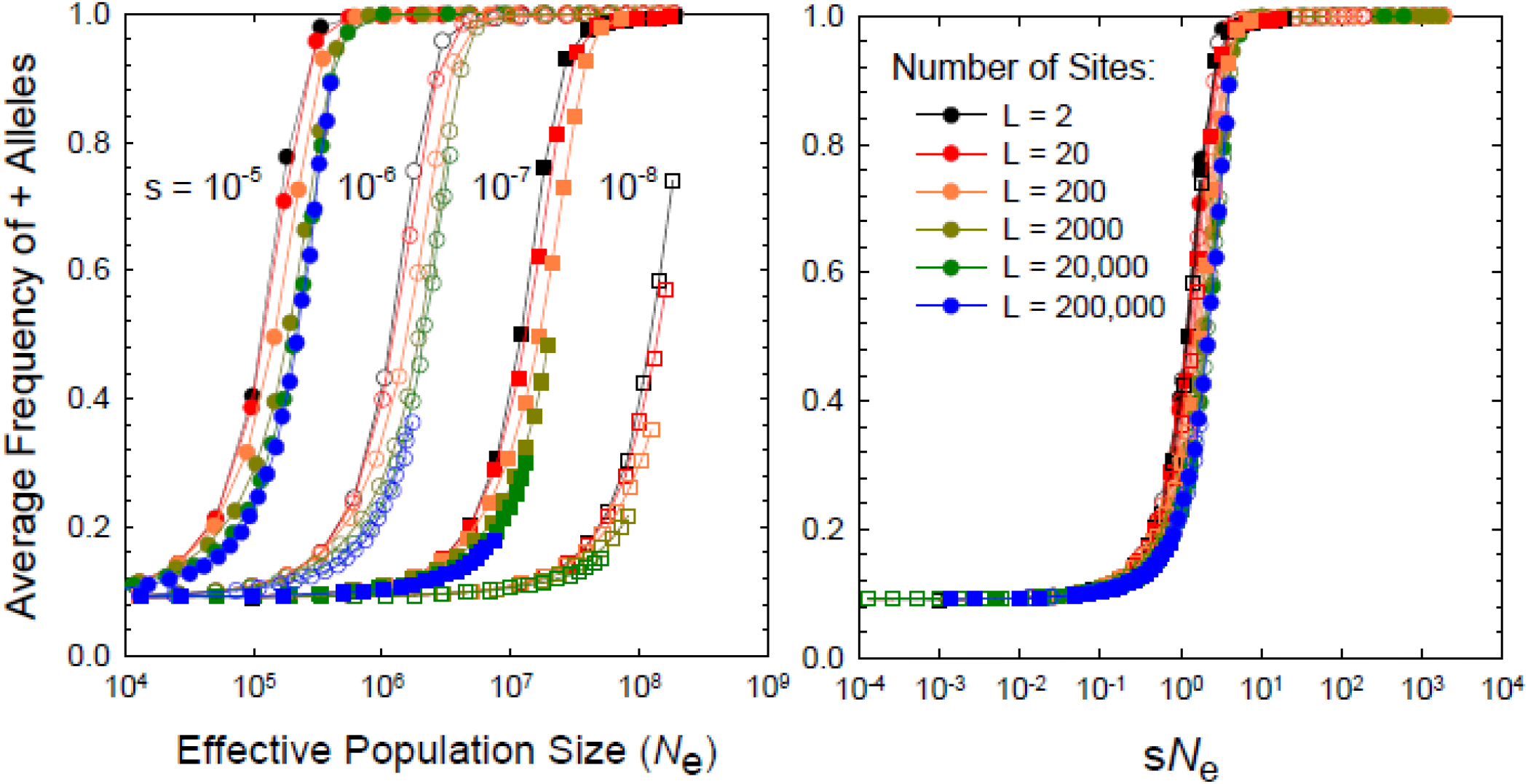
Response of the mean frequency of the beneficial allele as a function of the variance effective population size *(N*_*e*_ as determined from the long-term average diversity of variation at linked neutral sites) and of the product *N*_*e*_*s*, over a six order-of-magnitude range for *L* (the number of linked sites) and a four order-of-magnitude range of *s*.

Finally, note that the preceding expressions also yield descriptions of the expected standing levels of variation for quantitative traits under persistent directional selection (in this case an exponential fitness function) with reversible mutation, a problem of long-standing interest in quantitative genetics (Walsh and Lynch 2018). For example, simplifying from Equations 2a,b, assuming unbiased mutation (*β* = 1), the average genetic variance for a trait with *L* equivalent loci with average squared allelic effect *E*(*a*^2^) is

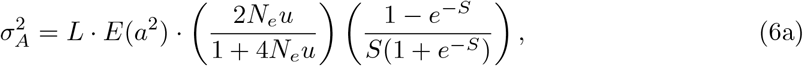

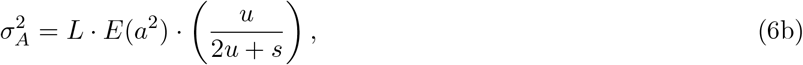

for *u* = *u*_10_ = *u*_01_, and *S <* 4 and *S >* 4, respectively. These expressions show that under selection, the genetic variance reaches a maximum at the point where 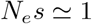, where the power of drift and selection are essentially equivalent (Supplemental Figure 1). The genetic variance initially grows with *N* owing to the increase in number of mutating individuals in the population, but beyond the peak, the deterministic force of selection overwhelms drift.

Even in the case of neutrality, there is a natural upper bound on the genetic variance, owing to the finite number of effects per nucleotide site (here assumed to be two), with the neutral variance in the case of *β* = 1 being simply proportional to 2*Nu/*(1 + 4*Nu*). Although it might be assumed that increased efficiency of selection (higher *Ns*) will always reduce standing levels of variation, in fact when mutation is biased in the opposite direction of selection, the genetic variance increases at an accelerating rate with *S* up to ≃ 4. This is because the conflict between mutation towards − alleles and selection towards + alleles pulls the latter towards more intermediate frequencies.

### Sites with two effects

Having arrived at a reasonable understanding of the factors determining the mean and variance of traits in the simplest case of *L* equivalent sites, we now explore the consequences of sites with variable fitness effects, starting with the case of just two site types to help illuminate the general complexities that arise. Some prior work has been done in this area (e.g., Johnson and Barton 2002; Desai and Fisher 2007; Pénisson et al. 2017; Jain 2019), but again in the context of an effectively infinite population size and an infinite-sites model. Here, we assume that the two site types have identical mutational features, while allowing for different site numbers.

Results described in the preceding section show that when linked sites have single effects, there is a smooth gradient in the expected frequency of favorable alleles with increasing population size. For any particular *s*, the gradient with *N* becomes increasingly shallow with increasing numbers of linked loci, owing to enhanced levels of selective interference, which causes an increasing fractional reduction in the effective population size. This gradient becomes steeper and is almost independent of *L* when reformulated as a function of *N*_*e*_ rather than *N*.

However, when sites with two effects contribute to the expression of a trait, a qualitative shift in the response of the mean phenotype to *N*_*e*_ is expected, as the sites with larger fitness effects will make a transition to high frequencies at a lower *N*_*e*_ than those with small effects. Moreover, a shift in the scaling of the fixation *N*_*e*_ with respect to *N* can be anticipated owing to the fact that once *N* is high enough to enable all major-effect sites to approach fixation for + alleles, these sites no longer contribute much to selective interference.

Consider, for example, a trait having an underlying additive-genetic basis with two types of sites: *L*_*M*_ sites with major phenotypic and fitness effects *a*_*M*_ and *s*_*M*_, and *L*_*m*_ sites with minor effects *a*_*m*_ and *s*_*m*_. The mean genotypic value is then

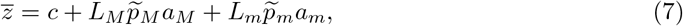

where 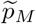 and 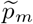 denote the mean frequencies of the + alleles at the major and minor loci, and *c* is an arbitrary baseline constant. If *s*_*M*_ is substantially larger than *s*_*m*_, beneficial alleles at the major-effect sites will achieve near fixation at relatively low *N*_*e*_, before the minor-effect sites begin to respond to selection.

An example is shown in Figure 5, where the response of mean performance to the effective population size in the single-effects case (equivalent to 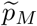) is compared to that for cases in which there are ten-fold additional minor-effect sites for each major-effect site, each with ten-fold lower selection coefficients, i.e., *L*_*m*_ = 10*L*_*M*_, and *s*_*M*_ = 10*s*_*m*_. In this figure, *N*_*e*_ is the variance effective size inferred by the average level of nucleotide diversity at linked neutral markers, as this would typically be the measure used in a population-genetics analysis. Assuming the phenotypic effects are proportional to the selection coefficients, mean performance in the two-effects case is defined by Equation 7, normalized by 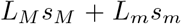 to give a maximum performance of 1.0 when all sites are fixed for + alleles. In this example, with 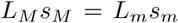, half of the maximum performance is determined by each class of sites. With a ten-fold difference in *s* between classes, as *N*_*e*_ increases to the point at which 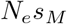 exceeds 1.0, a shoulder appears in the response profile because most major-effect loci are near fixation for + alleles, whereas the minor-effect loci do not start to significantly respond to selection until 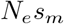 approaches 1. As a consequence, whereas the dynamic range of performance extends over just one order of magnitude of *N*_*e*_ under the single-effects case, the gradient extends for two orders of magnitude of *N*_*e*_ when substantial numbers of minor-effect sites are present.

**Figure 5.**
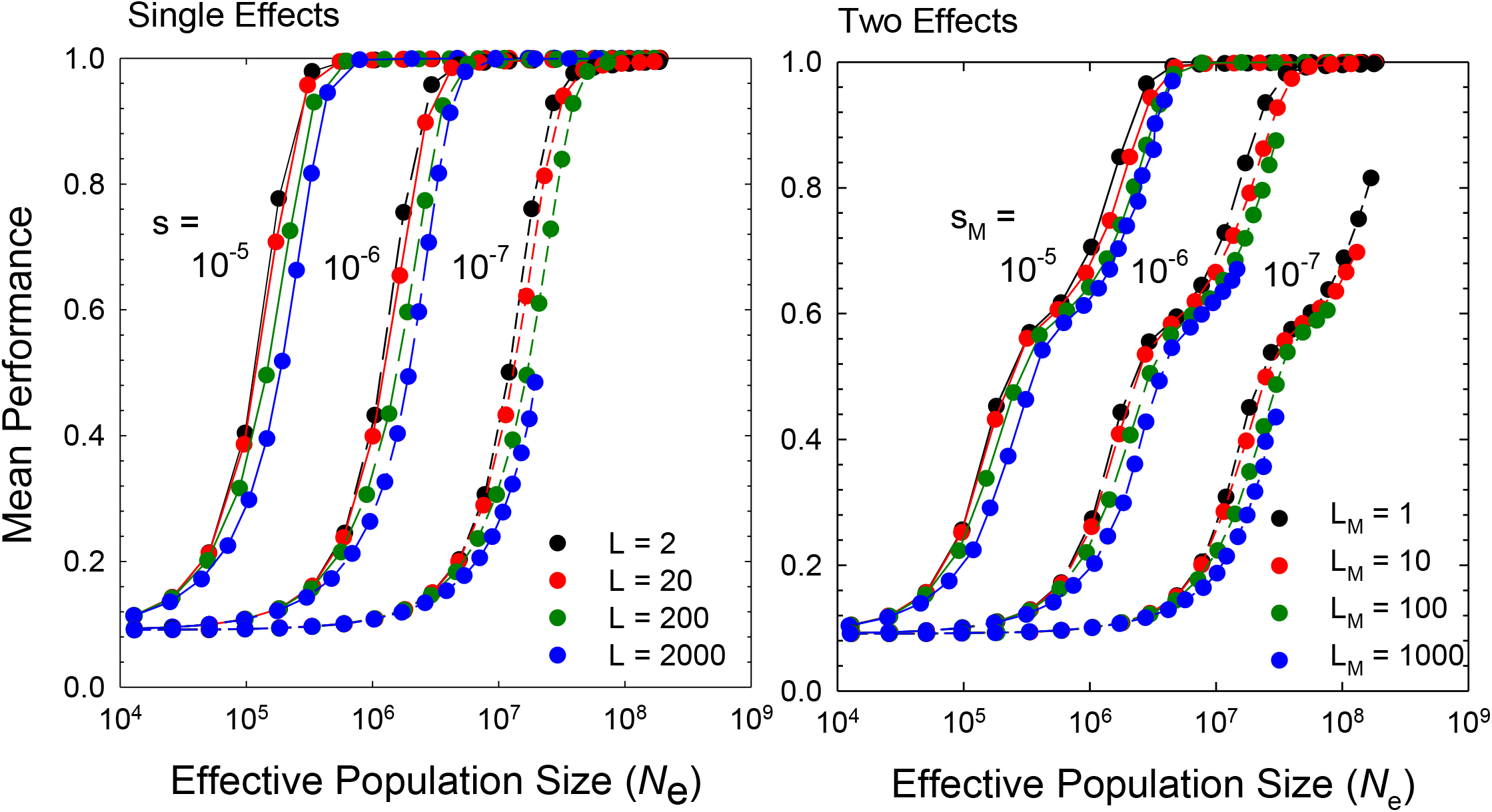
Response of mean performance to *N*_*e*_ (as determined from the nucleotide diversity at a linked neutral site) for the case of one- and two-effect models. In the two-effects case, the minor-effect loci have one-tenth the selection coefficient as that for the major-effect sites (*s*_*M*_ = 10*s*_*m*_) but are 10× more abundant (*L*_*m*_ = 10*L*_*M*_), such that the total potential selective load is the same for both types of sites, i.e., 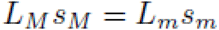.

All of the issues raised in the preceding section on selective interference effects between linked loci apply here, except that there is an asymmetry in the degree of selective interference that depends on the relative abundance of the two site types. Most notably, from the standpoint of major-effect sites, there is a remarkable simplicity with respect to the interference caused by minor-effect loci (Supplemental Text). The interference of a single minor-effect locus imposed on major-effect loci is equivalent to the influence of (*s*_*m*_*/s*_*M*_)^2^ of the latter (Figure 6). This yields an overall level of interference operating on major-effect loci equivalent to that resulting from

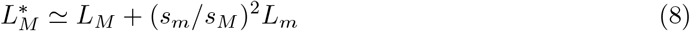

sites in the single-effects model. In other words, from the standpoint of a major-effect site, if *s*_*m*_*/s*_*M*_ = 1*/*10, the addition of 100 linked minor-effect sites is required to shift the effective amount of interference from that associated with *L*_*M*_ to *L*_*M*_ + 1 major-effect sites. All of the machinery just introduced for estimating the behavior of linked major-effect loci can then still be relied upon by substituting 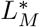 for *L*. This scaling with the squared effect of the selection coefficient can be roughly understood by noting that the probability of an establishment of beneficial mutation of effect *s* is also proportional to *s*.

**Figure 6.**
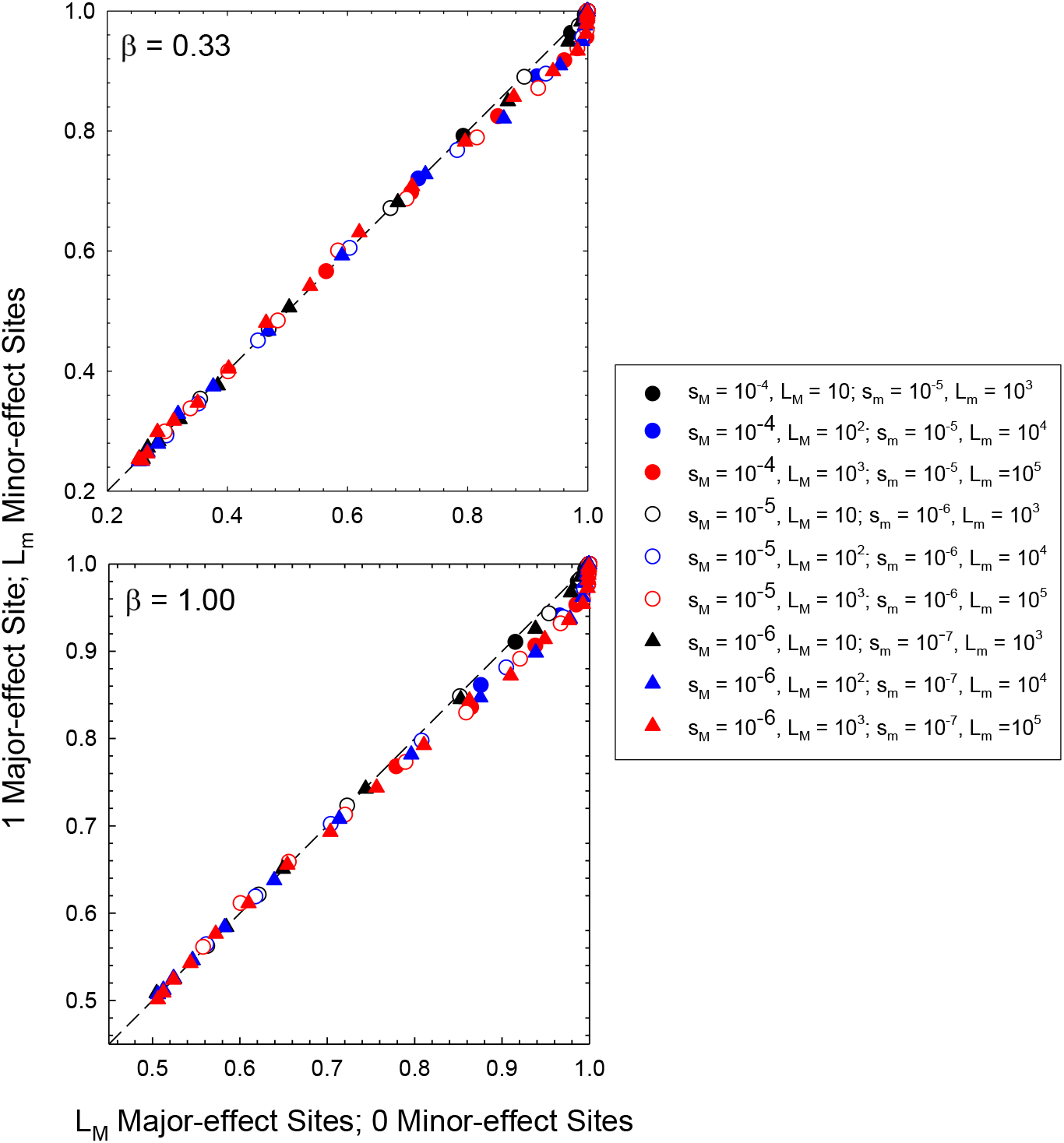
Demonstration that the effects of selective interference of a minor-effect site on the equilibrium frequencies of beneficial alleles at major-effect sites is equivalent to adding (*s*_*m*_*/s*_*M*_)^2^ major-effect sites to the genome. ‘fhe values on the *x* axis refer to results under the single-effects model with various numbers of loci with major effects *(L*_*M*_ in the insets) and no background minor-effect sites. The values on the *y* axis refer to results when a single major-effect site is surrounded by *L*_*m*_ minor-effect sites. Each point denotes the equilibrium mean frequency of + alleles under both conditions. In all cases here, *s*_*m*_*/s*_*M*_ = 0.1, and the prediction is that (*s*_*M*_*/s*_*m*_)^2^ = 100 minor sites have the same influence on the equilibrium + major allele frequency as the addition of one more major-effect site. Computer simulation results are given for 21 dill rent population sizes for each set of conditions. For every set of points on the *x* axis witl1 *L*_*M*_ major-effects sites alone, there is a parallel set of points on the *y* axis with 100*L*_*M*_ minor-effect sites but just a single actual major-effect site.

More generally, the overall fixation effective size of the population (which must be the same for both site types within linkage blocks) is dominated by the sites for which the product 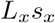 is highest, provided the population size is not so great that the categories in question are in near selection-mutation balance (i.e., no longer influenced by the vagaries of random genetic drift or selective interference). This will now be illustrated by considering three alternative domains of relative values of 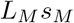 and 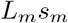 (Figure 7).

**Figure 7.**
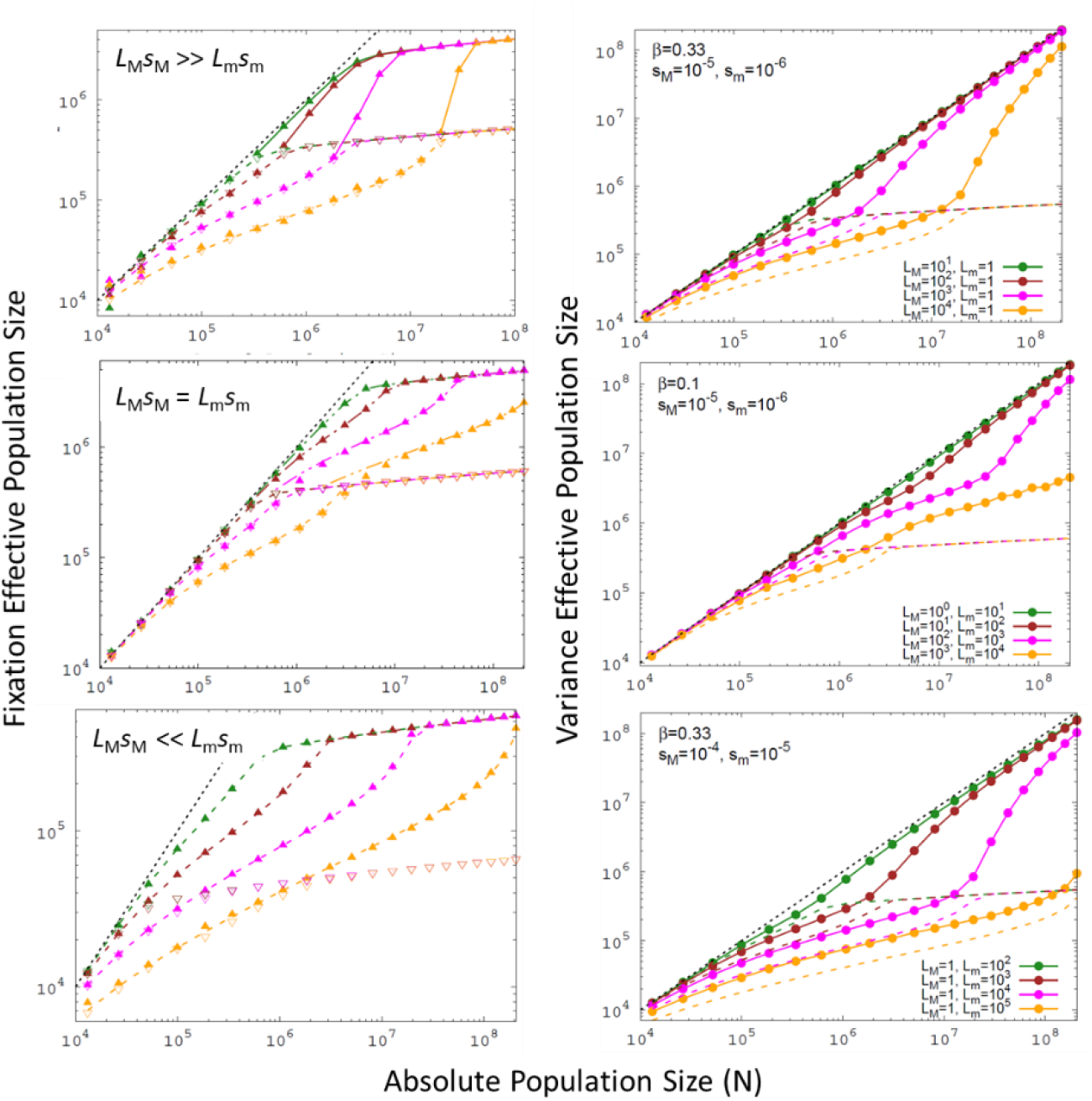
Relationship between fixation (left) and variance (right) effective population sizes to absolute population sizes, for three relative conditions involving 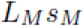 and 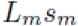, Left) Open and closed points denote results for major- and minor-effect sites, substituting in the mean frequencies of allelic types obtained by computer simulations into Equation 3 and multiplying by *N*. Dashed lines denote results obtained with Equation 3, using: *L* = *L*_*M*_ and *s* = *s*_*M*_ in the top panel; *L* = *L*_*M*_ and *s* = *s*_*M*_ in the lower left and *L* = *L*_*m*_ and *s* = *s*_*m*_ in the upper right in the middle panel; and *L* = *L*_*m*_ and *s* = *s*_*m*_ in the bottom panel. Solid lines simply join the connecting points. Note that regions of the plots where the fixation *N*_*e*_ levels off are within the pseudo-Ne domain, where sites have nonzero equilibrium frequencies of deleterious alleles defined by selection-mutation balance. **Right)** Solid points give the variance effective population sizes obtained from simulations of linked neutral sites and factoring out the mutation rate from the mean observed neutral heterozygosity to obtain *N*_*e*_. Dashed lines are taken from the right panels to compare the variance and fixation effective sizes. In all cases, the black dashed line denotes *N*_*e*_ = *N*.

First, for the extreme situation in which 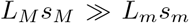, mutations at the minor-effect sites are subject to strong hitchhiking effects associated with the major-effect backgrounds upon which they arise. However, the major-effect sites behave in accordance with the predictions from the single-effects model, as they experience essentially no interference from minor-effect sites. With complete linkage, the behavior of the major-effect sites then dictates the *N*_*e*_ for the entire linkage block. As noted above for the single-effect model, the fixation effective *N*_*e*_ for major-effect sites steadily increases with *N*, to a degree that depends on *L*_*M*_, but upon reaching a critical *N* levels off as all such sites are close to pure mutation-selection balance, and then enters into a pseudo-*N*_*e*_ domain.

This pseudo-*N*_*e*_ domain is purely a mathematical feature of the use of Equation 3 to define the fixation *N*_*e*_. As *N* reaches high enough levels that 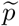 approximates the level expected under pure selection-mutation balance, absolute fixation of + alleles never occurs, although genealogical fixation does. As a consequence, the fixation *N*_*e*_ levels off, as *ϕ*_max_ (Equation 5a) declines. Although this pseudo *N*_*e*_ is not a reflection of the actual *N*_*e*_ in the large-*N* domain, its deployment in the preceding mathematical expressions is required to obtain an acceptable overall expression for 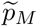. The behavior of the variance *N*_*e*_, shown in the right panels of Figure 7, yields some insight into the stochastic features of the population, as it qualitatively tracks the behavior of the fixation *N*_*e*_ (outside of the pseudo-*N*_*e*_ domain), although overestimating the latter. The variance *N*_*e*_ always starts out as *N*_*e*_ = *N* at 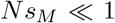, is reduced relative to *N* at intermediate *N* by selective interference, and then asymptotically returns to *N*_*e*_ = *N* for 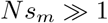.

In the example shown, because *s*_*m*_ = *s*_*M*_ */*10, the minor-effect sites do not begin to respond to selection until *N* is an order of magnitude beyond the point at which the major-effect sites are subject to selection. At this point, the governing fixation *N*_*e*_ is reflected in the behavior of the minor-effect sites, until they too enter their pseudo-*N*_*e*_ domain at very large *N*. We have not been able to achieve a fully mathematical description of this transitional behavior in the minor-effect sites in this limiting case of 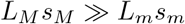, but an intuitive understanding of the processes involved can be understood as follows.

One might expect the fixation *N*_*e*_ for minor-effect alleles to increase to *N* once the deterministic regime for major effects has been reached, as in this case there is a just a single minor-effect site. However, from the behavior of the variance *N*_*e*_, it can be seen that the system does not return to *N*_*e*_ = *N* until *N* is well beyond the point of entry of the major-effect sites into the deterministic regime. Selective interference from the major-effect sites still occurs (to a degree that increases with *L*_*M*_), owing to the background variation among individuals with respect to major-effect alleles. For example, at *N* ≃ 10^6^, in this particular set of simulations, the mutation rate to deleterious alleles ≃ 10^*−*8^, so with *s*_*M*_ = 10^*−*5^, the equilibrium frequency of deleterious alleles ≃ 10^*−*3^. With *L*_*M*_ = 10^3^, there is then an average of 1.0 deleterious major-effect mutations per individual, and as the distribution among individuals is expected to be Poisson, ∼ 37% of individuals will be free of deleterious major-effect mutations. Only in this subset of individuals are beneficial minor-effect mutations able to progress towards fixation, as all lineages containing major-effect deleterious alleles will be subject to purging from the population (unless a reversion mutation is acquired), and even then on a large *L*_*M*_ background, some of these can become victims of subsequently arriving linked major-effect deleterious mutations. Thus, if *L*_*M*_ is sufficiently large, trapping of beneficial minor-effect mutations can impose selective interference effects even in the deterministic regime for major-effect sites.

We next consider the situation in which 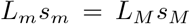, such that there is an inverse relationship between the number and selective effects associated with the two site types. In this case, at sufficiently small *N* such that the minor-effect sites are effectively neutral, the fixation effective population size is a function of 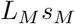. A departure between the estimates of *ϕ* using major-vs. minor-effect sites only arises at larger *N* as the major-effect sites enter the pseudo-*N*_*e*_ domain (*ϕ*_max_ *<* 1), while the minor-effect sites remain under the stochastic effects of drift. At this point, the fixation *N*_*e*_ of the minor-effects sites is primarily a function of 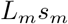, until they themselves enter their pseudo-*N*_*e*_ domain. This kind of domain shift will become more blurred as *s*_*m*_ → *s*_*M*_, in the limit becoming equivalent to the single-effects model with *L* = *L*_*M*_ + *L*_*m*_.

Finally, for the case in which the minor-effect sites greatly outnumber those for major effects, such that 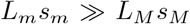, the former are largely unaffected by interference from the major-effect sites, rendering the behavior of the minor-effect sites very close to that observed in the single-effects situation as defined by *L*_*m*_ and *s*_*m*_. In this case, the behavior of the major-effect sites is also essentially defined by the minor-effect loci, as 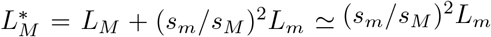. In both this and the prior case, background trapping of minor-effect alleles plays a negligible role in the pseudo-*N*_*e*_ domain for the major-effect sites because much to most of the background variation is associated with the minor-effect sites.

### Generalization to multiple site types

With sites with additional effects, one can anticipate an extension of the features noted above. From the standpoint of the major-effect loci, the preceding logic can be extended to an arbitrary number of effects, yielding an interference effective number of major sites equivalent to

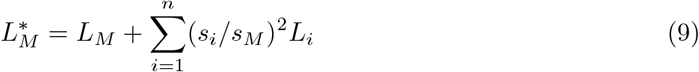

where *L*_*i*_ is the number loci with effect size *s*_*i*_ *< s*_*M*_, and *n* is the number of effect classes. For many polygenic traits, the distribution of site types may be nearly continuous in form, in which case this expression could be replaced by an integration over the full spectrum of site types.

Examples are given in Figure 8 for the case of three effects with an inverse relationship between site numbers and effects, such that 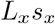 is constant, and with ten-fold differences in *s*_*x*_ between site types. In this situation, the summed effects of sites within each of three classes contribute equally to the total performance of the trait (assuming additive phenotypic effects across sites). As the population size declines, the site types with smaller *s* progressively accumulate deleterious mutations, yielding a gradient for mean total performance that is nearly continuous over several orders of magnitude of *N*. Increasing the number of sites within linkage blocks (while keeping the ratios of numbers of site types constant) has a greater effect on the sites with small effects, reducing the steepness of the performance gradient. The precise form of the scaling would be altered with different distributions of effects, which would shift the relative contributions of the three types to mean performance.

**Figure 8.**
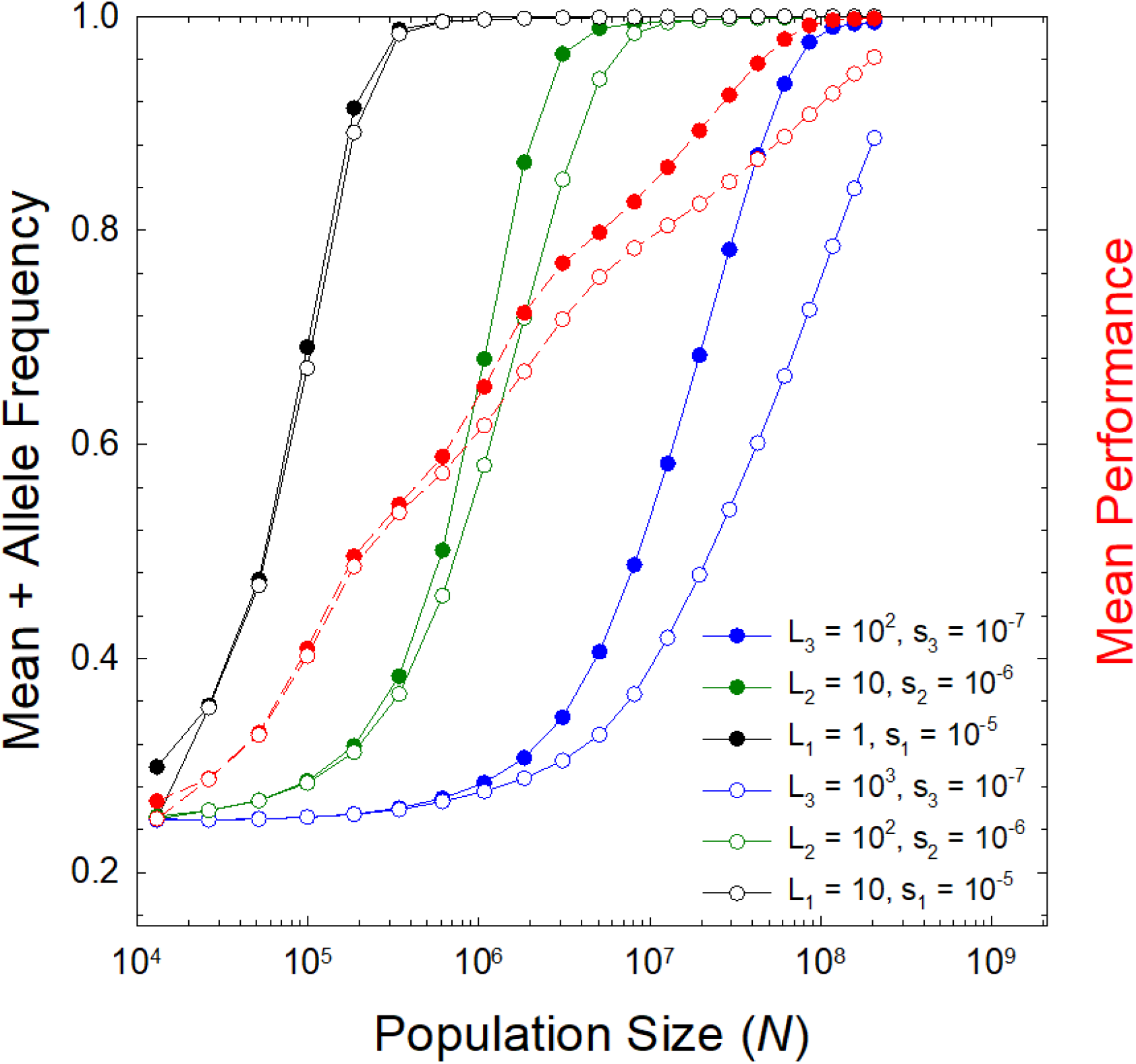
Response of mean + allele frequencies to population size (*N*) for the case of the three site types, with an inverse relationship between the number of sites and the selective effects within a class, such that 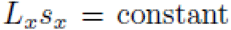, with a ratio of 1:10:100 for major:medium:minor-effect sites. The mutation bias towards + alleles is set to *β* = 0.33. Two situations are shown, with the major site type being present in one (solid points) or ten (open points) copies. Mean performance is obtained by extension of Equation 7 to three site types, normalized by the value expected when all sites are fixed for + alleles.

## DISCUSSION

The primary motivation for this work is the idea that quantitative traits under persistent directional selection of the same form in different phylogenetic lineages should exhibit gradients in mean phenotypes associated with differences in effective population sizes. Such an expectation arises for the simple reason that *N*_*e*_ dictates the efficiency of natural selection, and should hold generally provided that a significant fraction of mutations with phenotypic effects have selection coefficients within the lower and upper bounds of 1*/N*_*e*_ across phylogenetic lineages. The fact that *N*_*e*_ ranges from 10^4^ to 10^9^ among phylogenetic lineages, scaling negatively with the ∼ 0.2 power of body mass (Lynch and Trickovic 2020), indicates that the complete absence of the effects of drift on mean phenotypes requires an absence of mutations with fitness effects in the range of 10^*−*9^ to 10^*−*4^, which seems highly implausible.

Prior theoretical work provided a framework for considering the fundamental population-genetic processes influencing the generation of such gradients, including the effects of linkage block size (an analog of the level of recombination), but under the restriction that all mutations have comparable effects on phenotypes and fitness (Lynch 2020). Here, we have provided more general mathematical approximations for the single-effects model, and used these to further understand the more biologically realistic situation in which genomic sites have different effects. Ultimately, we would like to make statements on the quantitative scaling of mean phenotypes with *N*_*e*_ based on first principles, but this will require detailed information on the distribution of mutational effects summarized over different site types. Unfortunately, the fraction of this distribution that is of most relevance resides within the 1*/N*_*e*_ bounds noted above. Although likely quite abundant, mutations with such small effects are highly impenetrable to direct enumeration (Walsh and Lynch 2018; Lynch and Ho 2020). For now, we at least have a framework within which to derive predictions under specified genomic and population-genetic conditions.

For example, although a fully general description of the steady-state distribution of mean phenotypes under a variable-effects model remains to be developed, the preceding results provide the basis for a heuristic argument as to how mean phenotypes under persistent directional selection should scale with *N*_*e*_. To clarify the main points, the following qualitative discussion relies on order-of-magnitude arguments, and starts with a simple additive genotype-to-phenotype mapping,

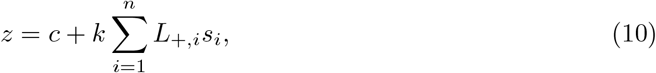

such that the expected trait value is a linear function of the number of plus alleles at each of *n* types of sites, each weighted by the selective advantage, with *c* and *k* being arbitrary constants. We wish to determine how the mean genoytpic value 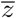 scales with *N*_*e*_.

As noted above, *N*_*e*_ will be largely governed by the sites with the largest fitness effects, unless there is a much stronger than exponential increase in the numbers of sites with diminishing effects. Supposing the sites with the strongest effects have *s* = 10^*−*5^, then below *N*_*e*_ = 10^4^, all such sites will have expected + allele frequencies at the neutral expectations defined by the level of mutation bias. Above *N*_*e*_ = 10^6^, all such sites will be essentially fixed for + alleles, with most of the gradient residing in the vicinity of *N*_*e*_ ≃ 10^5^. Likewise, sites with *s* on the order of 10^*−*6^ will exhibit a gradient in the vicinity of *N*_*e*_ = 10^6^, with + alleles just starting to accumulate at *N*_*e*_ ≃ 10^5^ and becoming essentially fixed at *N*_*e*_ ≃ 10^7^. The same argument applies to sites with all lower-order effects, with each order-of-magnitude effect exhibiting a gradient roughly corresponding to where the prior and subsequent ones exhibit maximum responses to *N*_*e*_.

The precise form of the gradient of 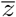 will depend on the relative incidences of site types and on the form of the genotype-to-phenotype map. For example, for the linear mapping in Equation 10, if the number of sites of type *i* is inversely proportional to *s*_*i*_ (i.e., an essentially exponential distribution of site types), then the contribution of each site type to total performance will be equal over all site types, as in Figure 8, and there will be a continuous gradient over *N*_*e*_ in the range of 1*/s*_*max*_ to 1*/s*_*min*_. (The small wobbles in the gradient shown in Figure 8 would become essentially invisible with the inclusion of more fine-grained effects). Deviations from an exponential distribution of site types would alter the gradient accordingly, as would a different weighting scheme for the genotype-phenotype map. For example, if there was a paucity of intermediate-effect sites, there would be a shoulder in the response to *N*_*e*_, as nearly all large-effect sites would become fixed before *N*_*e*_ reaches a high enough level for small-effect sites to respond to selection.

While not a formal mathematical statement, this heuristic argument provides a roadmap for thinking about how phenotypic gradients should scale with effective population sizes for traits under similar forms of directional selection across species. If, for example, there was an exponential distribution of sites with different fitness effects over five orders of magnitude of *s*, a gradient in performance would be expected over five orders of magnitude of *N*_*e*_. The actual “power-law” scaling would depend on the scale upon which performance is measured and on the genotype-phenotype map. For a multiplicative mapping function, on a logarithmic scale (the usual procedure in studies of allometry), the expected slope might approach +1, but could be shallower in the case of large linkage blocks, which would enhance the level of selective interference. In contrast, the linear mapping function used above might lead to nonlinear allometric scaling, depending on the distribution of site types.

There is considerable room for more theoretical work in this area. For example, as a surrogate for the level of recombination, we have relied on the concept of a linkage block (Good et al. 2014), which greatly facilitates computational study in the domain of large *L*, and also eases a number of aspects of the mathematical analysis. Although we do not expect qualitative changes in the conclusions to result from a more fully implemented recombinational model, work of this nature is desirable. Most notably, we have focused on a single fitness function (albeit a common one used in studies of deleterious-mutation accumulation), the exponential (or multiplicative) model, wherein there are no epistatic effects of mutations, as each additional deleterious mutation reduces fitness by a fractional amount *s* regardless of the genetic background. Variants of this model have been invoked to explain how drift barriers may influence the phylogenetic distribution of mutation rates (Lynch 2011; Lynch et al. 2016) and maximum growth rates (Lynch et al. 2022), both of which are plausibly under persistent directional selection in most lineages. Other types of traits that might be explored in this regard are cell biochemical and/or physiological processes shared across the Tree of Life.

Future exploration will need to consider Gaussian and mesa fitness functions, which do introduce epistatic effects. The mesa fitness function (with a plateau) imposes pure directional selection with diminishing fitness increments as the trait approaches the asymptotic optimum, whereas the Gaussian fitness function provides a setting in which traits can be under stabilizing selection for an intermediate optimum. In the limit, as the optimum falls far out of the range of obtainable phenotypes, the Gaussian fitness function converges on the exponential model used herein.

Notably, although a substantial body of work in evolutionary quantitative genetics has been developed under the assumption of a Gaussian fitness function, and many evolutionary biologists operate under the assumption that such stabilizing selection is pervasive, a broad survey of estimated fitness functions raises questions about the generality of this model and certainly leaves open the possibility that persistent directional selection is a common force (Kingsolver and Diamond 2011). In the field of evolutionary ecology, arguments for suboptimal performance traditionally invoke limitations owing to constraints / tradeoffs between traits, which are typically assumed but seldom verified empirically. Here, we have shown that persistent under-performance can be expected whenever a significant fraction of genomic sites contributing to a trait harbor preferred alleles with small selective advantages, and that this effect will become more pronounced when mutation is biased in the direction of deleterious alleles. Some progress has been made on the study of the expected evolution of mean phenotypes under these alternative models (Charlesworth 2013b; Lynch 2018, 2020), but again under the assumption of mutations with fixed effects. Our results show that substantially different conclusions may arise under more biologically realistic scenarios when genomic sites are variable in their effects.

## Acknowledgments

This research was supported by the Multidisciplinary University Research Initiative awards W911NF-09-1-0444 and W911NF-09-1-0444 from the US Army Research Office, National Institutes of Health award R35-GM122566-01, National Science Foundation awards DBI-2119963, DEB-1927159, and MCB-1518060, and Moore and Simons Foundations Grant 735927.

## Data availability

The authors affirm that all data necessary for confirming the conclusions presented in the article are represented fully within the article and figures. The C++ code for the simulation data can be found on the GitHub website (https://github.com/ArchanaDevi8474/ThreeEffectsSimulationCode).

## SUPPLEMENTAL MATERIAL

### Reduction in *N*_*e*_ by selective interference: single-effects case

Observations reported in the text justify the use of a correction factor *ϕ* = *N*_*e*_*/N* to transform *N* into a *fixation effective population size* relevant to the evolution of mean allele frequencies, validation of this approach required the use of estimates of *ϕ* derived by computer simulations. For more practical applications, we require an expression for *ϕ* from first principles. A heuristic approximation can be obtained by considering the number (*I*) of competing mutations that a mutation destined to fixation must contend with during its sojourn through the population. Gerrish and Lenski (1998) and Campos and Wahl (2009, 2010) used such an approach to evaluate the number of newly arising mutations with advantages exceeding that of a target mutation, under the assumption of an exponential distribution of mutational fitness effects, but here all newly arising beneficial mutations have identical effects.

We start with the Li-Bulmer equation for the expected frequency of a beneficial allele under sequential fixations

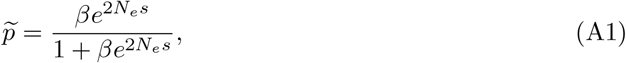

where *β* = *u*_01_*/u*_10_ is the ratio of mutational pressure towards the beneficial relative to the deleterious allele, *s* is the selective advantage of the beneficial allele, and *N*_*e*_ is the effective population size. Rearrangement of Equation A1 leads to

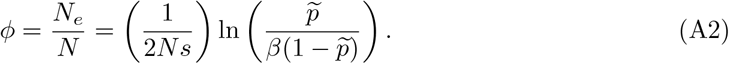

Although Equation A1 implies that 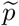 asymptotically approaches 1.0 as the population size approaches infinity, in this extreme, deleterious alleles will actually be maintained at a low frequency by mutation-selection balance, such that

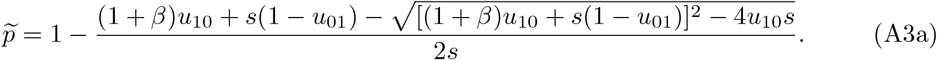

Provided the strength of selection exceeds *u*_10_,

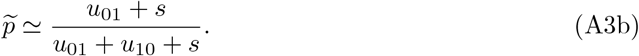

Substituting Equation A3b into A2 then yields an upper bound to *ϕ*,

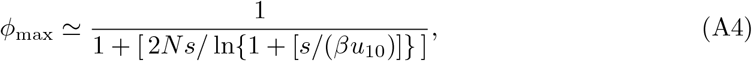

where a slight modification has been made by adding 1 to the denominator to account for the fact that *ϕ*_max_ must asymptotically approach 1 as *N* → 1 declines (as this eliminates interference). Consistent with this expression, the simulation results show that once *Ns* exceeds 10, *ϕ* becomes inversely proportional to *Ns*, and only weakly dependent on the composite selection-mutation parameters subsumed into *s/*(*βu*_10_) (Figure 4). To allow for the further depressive effects of selective interference from linked mutations, we use

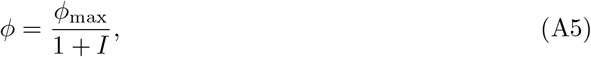

where it is assumed that when combined with *I* interfering mutations, a target mutation destined to fixation in the absence of interference has its probability of fixation reduced by factor 1*/*(1+*I*). This approach ignores the possibility that mutations interfering with the fixation of a focal beneficial mutation can also interfere with themselves.

We now proceed towards the development of an estimator for *I*, progressively accounting for the number of potentially interfering mutations arising during a focal mutation’s sojourn through the population, along with the magnitude of the effect per interfering mutation. Let *τ* be the mean time to fixation of a beneficial allele in the absence of competition from other segregrating mutations. During this period, additional beneficial mutations will arise in individuals outside of the focal lineage at average rate *Lβu*_10_(1 − *p*^*^), where *p*^*^ denotes the expected mean frequency of + alleles. The average per-generation number of individuals in the target lineage is *N/*2 because the frequency of the lineage under consideration (assuming it does indeed fix) increases from essentially zero to one. Only a fraction *p*_*f*_ (*s*) of all newly arisen beneficial mutations are destined to fixation, and it is this subset that presents the most potential interference to the focal mutation. Moreover, the strength of selection operating on a mutation must be on the order of the magnitude of genetic drift or greater if it is to compete for fixation; and to account for this, we use the weighting term 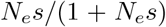, which asymptotically approaches zero as 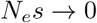 and 1.0 as 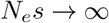.

Taking all of these factors into consideration, the expected number of competing mutations is then proportional to the product of terms,

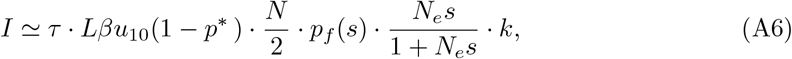

where

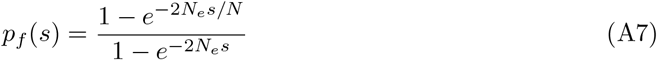

is the probability of fixation of a newly arisen mutation with fitness benefit *s* (Kimura 1983), and *k* is a correction term to account for the fact that only a fraction of newly arising mutations emerge in backgrounds with high enough fitness to compete with the target mutation.

Although *k* must depend on the distribution of fitness in the population, a rough starting point is *k* = 0.25, as any mutant haplotype destined to fixation by positive selection must almost certainly be in the upper half of the distribution, and to be a successful competitor, any outside mutant clone must then be in the upper half of the upper half. Previous workers (Gerrish and Lenski 1998; Campos and Wahl 2009, 2010) have let *k* = 1, which applies if prior to the emergence of competing mutations, the population consists of just two haplotypes, one with and the other without the target mutation. In principle, a more rigorous approach might be possible if the form of the equilibrium distribution were known, but although this is often assumed to be Poisson, there are subtle and significant deviations from such behavior in numerous contexts (Gessler 1995; Goyal et al. 2012; Jain and John 2016), and *k* ≃ 0.25 will be shown to be a reasonable approximation for an equilibrium population. Note also that the matter of lineage contamination, i.e., the addition of secondary mutations to lineages en route to fixation (Pénisson et al. 2017) has been ignored here, as the population is in equilibrium, and all competing mutant lineages are presumably confronted with the same secondary-mutation issues; in effect, as *N* → ∞, all mutant lineages approach selection-mutation balance, rendering a near neutral situation with respect to lineage competition.

Aside from the inclusion of the factor *k*, our computation of *I* differs in several significant ways from prior applications. First, rather than approximating the fixation probability as 2*s*, we implement the full formulation for *p*_*f*_ (*s*), as the former yields inappropriate estimates when 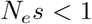 and ignores the fact that *N*_*e*_ *< N*, the very issue that we are exploring. Second, prior applications have not included the weighting term noted above, which also seems essential for 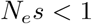, for which the magnitude of individual interference must be small. Third, the estimation of *ϕ* is quantitatively quite sensitive to the definition of *τ*, and whereas previous authors have used the deterministic approximation *τ* = 2 ln(*N*)*/s*, this can give wildly unrealistic values in certain domains of parameter space, including times in excess of the neutral expectation of *N*_*e*_ generations. Charlesworth (2020) provides a broad overview of estimators of *τ*, and offers a measure that explicitly accounts for the stochastic and deterministic phases of the process, but implementation of his method in Equation A6 consistently led to significant underestimates of *I*, whereas a derivation of Gale (1990), modified for haploids, was more suitable,

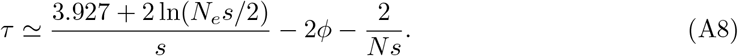

Note that the expressions of Gale (1990) and Charlesworth (2020) can yield negative estimates of *τ* when 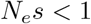, in which case we assumed the neutral expectation of *τ* = 2*N*_*e*_ generations. Finally, Equation A6 requires an expression for *p*^*^ (a concern given that *p*^*^ is the allele frequency that we are ultimately trying to determine), but here we utilize Equation A1 as a first-order approximation.

Equation A5 is a transcendental function, as several of terms entering *I* are complex functions of *N*_*e*_ = *ϕN* – the fixation probability and time, the weighting factor, and *p*^*^, although the equation can be solved by iteration. Despite the complexity of the underlying issues and the approximate nature of the derivation, Equation A5 generally yields estimates of *ϕ* that are within 30% (often considerably closer) of simulation results (Figure 4). Although this is not a fully satisfactory outcome, given that *φ* varies 10,000-fold over the full range of parameter space, this heuristic solution appears to capture the essence of the system, is an upgrade to the visual-fit interpretation of Lynch (2020), and provides insight into the regions of parameter space that merit further consideration. Major discrepancies appear to be restricted to very large linkage blocks (of order *L* = 10^6^) with *Ns* ≪ 1, where *ϕ* is underestimated up to two-fold, and with *Ns* in the range of 10 to 1000, where *ϕ* is overestimated up to five-fold. As can be seen in Figure 4, the primary determinant of *ϕ* is *Ns*, with *ϕ* only responding in a significant way after *Ns* exceeds a threshold value near 1 for small *L* and 0.01 for very large *L*. Not surprisingly, larger *L* leads to stronger interference, but in all cases, mutational bias (*β*) has a secondary effect.

A simpler approximate solution is obtainable under the assumption of 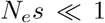, which allows the approximations *τ* ≃ 2*N*_*e*_ and 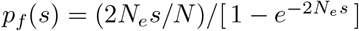, and thus to

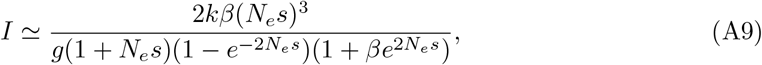

where *g* = *s/*(*Lu*_10_). Letting 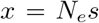 and referring back to Equation A5, this rearranges to another transcendental equation

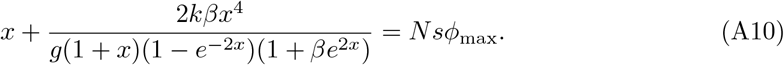

For small *x*, Taylor’s-series expansion of the left side of Equation A10 leads to

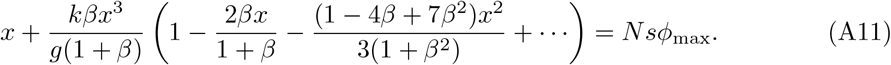

For small *L* and *u*_10_*/s* ≪ 1, the parameter *g* ≫ 1, and hence the above equation simplifies to *Nsϕ*_max_, and hence

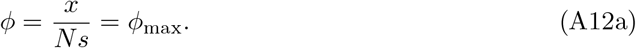

In the other limit, as *L* → ∞, the parameter *g* → 0, so the second term in the left side of Equation A11 dominates, leading to the approximation,

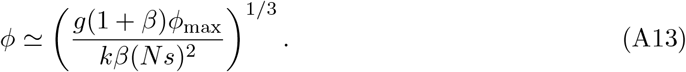

This expression yields scaling relationships that are fairly similar to those generated by less formal methods in Lynch (2020; Equation 11), which suggested *ϕ* to be an approximately inverse function of 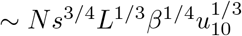 over the full domain of *Ns*, noting that here there are additional terms in *ϕ*_max_. Although Equation A13 can yield *ϕ >* 1 at small *Ns*, this can be accommodated by simply setting *ϕ* = 1 at this point.

For 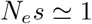, retaining the assumption of *τ* ≃ 2*N*_*e*_, using Equation A10, and performing a Taylor’s-series expansion around 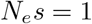 leads to

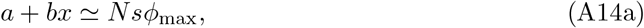

where

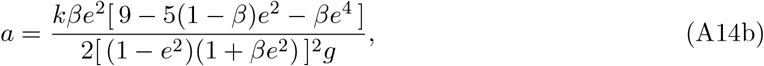

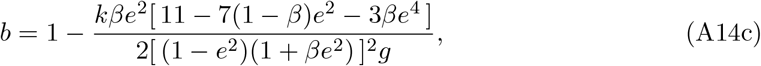

Equation A14a then yields

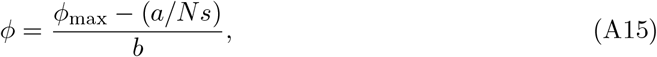

which converges to *ϕ* = *ϕ*_max_, as *g* → ∞.

Over the full range of parameter space for *L* = 10^6^, Equation A13 performs as well as (in some cases better than) the direct solution of the transcendental function (Figure 4). Equation A13 also performs reasonably well with *L* as low as 10^4^. For smaller *L*, Equation A13 tends to overestimate *ϕ*, whereas Equation A15 yields results that are nearly indistinguishable from the solution of the transcendental equation, which maps well to the simulation results. Note that in all evaluations here, we have retained the use of *k* = 0.25.

### Reduction in *N*_*e*_ by selective interference from minor-effect loci: two-effects case

A large body of literature in the concurrent mutation regime is based on a semi-deterministic approach where the bulk of the distribution of frequency classes follows a deterministic equation and the stochastic noise enters only at the nose of the distribution (Rouzine et al. 2008; Goyal et al. 2012; John and Jain 2015). However, these studies consider uniform sites with single mutational effects. Here, we try to understand the connection between the single-effects model and the two-effects case using the deterministic equation for the frequency class *P*_*i,j*_ containing *i* and *j* number of deleterious loci summed over all *L*_*M*_ and *L*_*m*_ major and minor-effect sites, respectively,

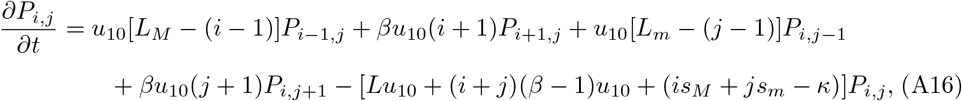

where *L* = *L*_*M*_ + *L*_*m*_ and 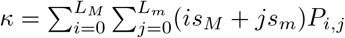.

Letting 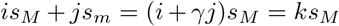, where 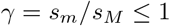, then by setting *k* = *i* + *γj*, the two-effects equation can be reduced to an effective one-effect equation with the frequency class *P*_*i,j*_ becoming *P*_*k*_ with associated selection *ks*_*M*_,

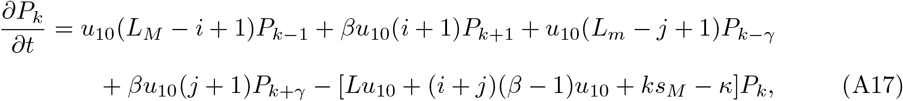

Dividing by *Lu*_10_ everywhere,

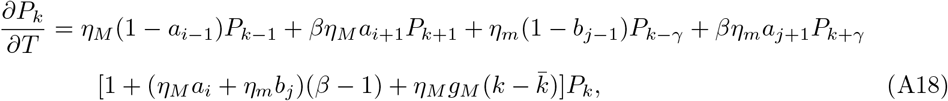

where 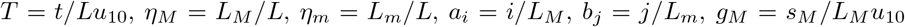, and 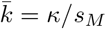.

When the joint distribution *P*_*i,j*_ is far from the edges, the approximations 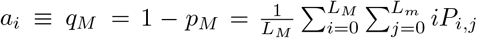 and 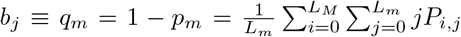.hold. Here, we follow the similar approach as in (Rouzine et al. 2008) and assume that the logarithm of the frequency is a smooth function, and use the approximation 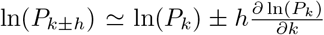. In the steady-state, the above equation then becomes

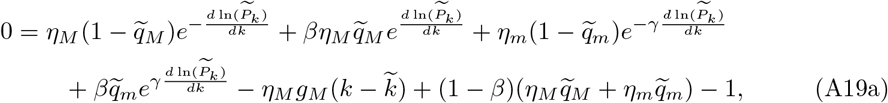

which after letting 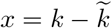 and 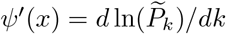 can be rewritten as

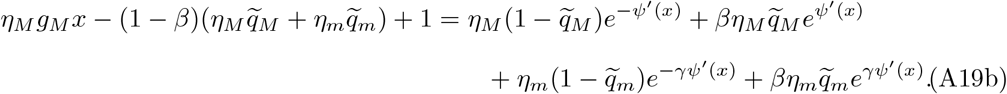

Although the above equation is not exactly solvable, by expanding the exponentials for small *ψ*′(*x*), we get

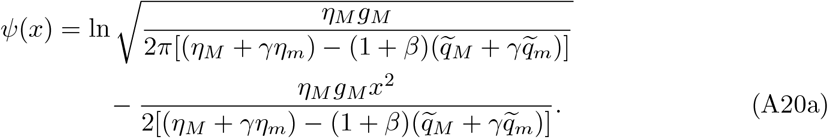

The first term on the right side is obtained from the normalization condition, ∫*e*^*ψ*(*x*)^*dx* = 1, assuming *x* is continuous. Equation A20a can be rewritten as

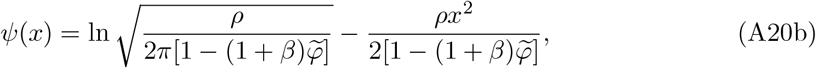

with 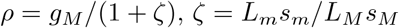, and 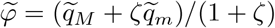. Since we have assumed the width of the distribution is large, this implies that the above calculation is valid for *ρ* ≪ 1. The above distribution yields a bivariate Gaussian distribution for *k* = *i* + *γj*,

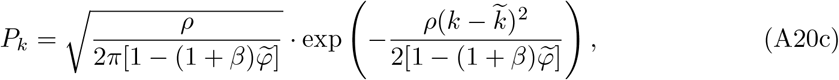

with mean 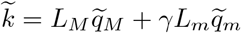 and variance 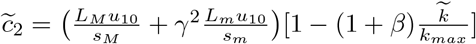 where *k*_*max*_ = *L*_*M*_ + *γL*_*m*_. Note that previous studies have also found this Gaussian distribution in the single-effects case both for infinite and finite loci models (Rouzine et al. 2008; Goyal et al. 2012; Jain and John 2016).

We can reproduce the single effects distribution by putting *s*_*M*_ = *s*_*m*_ or taking the number of major or minor loci to zero in (A20c). Here, the variance produced by the minor effects in the distribution of number of deleterious major loci is *γ*^2^ times smaller than that it can produce in the distribution of number of deleterious minor loci in the absence of major effects. In the presence of major effects, the variance caused by the minor loci in the distribution of number of deleterious major loci reduced by a factor of *γ*^2^, suggests that the influence of minor loci on the major effect is *γ*^2^ times its influence on itself. The major loci has the same influence on the major effects in the presence and absence of minor loci.

This variance in the distribution increases the interference effect which decreases the effective population size in a finite population due to linkage. We observe that the interference effect caused by the minor loci on the major loci follows the above discussion. We can also write the variance as 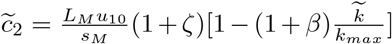 which indicates that the effective population size will be dominated by the major effects when *ζ* ≪ 1 or 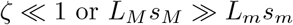 and by minor effects in the opposite regime.

---

**Supplemental Figure 1.**
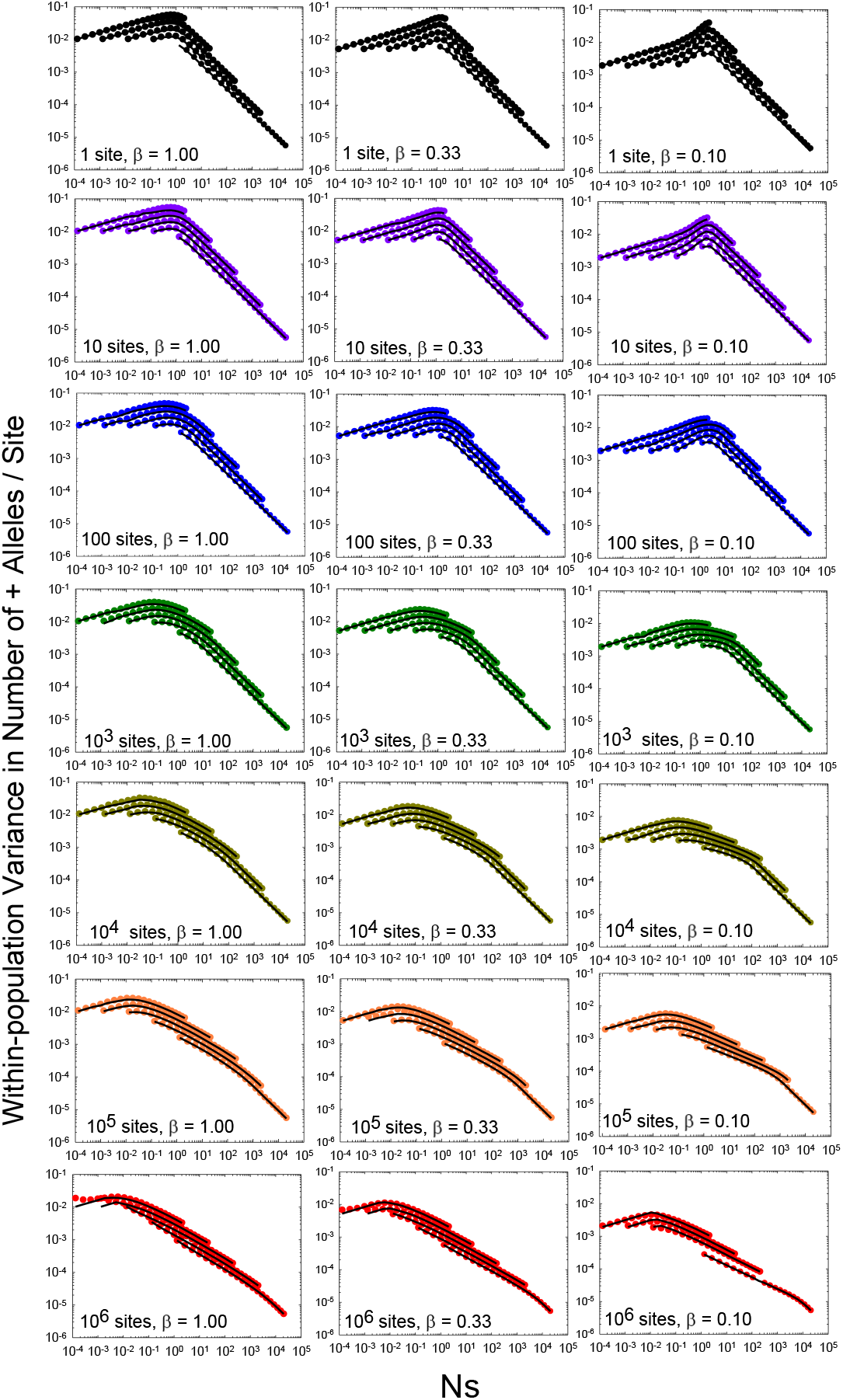
Scaling of the steady-state average single-locus within-population variance for presence of + alleles as a function of the scaled intensity of selection (*Ns*). Each row gives the results for three different levels of mutation bias, *β*= 1.0, 0.33, and 0.10. The case of free recombination (*L* = 1) is given in the first row, and then each row thereafter increases *L* by a factor of 10, up to *L* = 10^6^. Within each panel, there are five sets of results for different selection coefficients ranging in orders of magnitude from *s* = 10^−8^ (top leftmost curves) to *s* = 10^−4^ (bottom rightmost curves). The dots denote simulated data, whereas the black lines give the theoretical predictions based on Equations 2a,b.

## Literature Cited

Bulmer, M. G. 1985. The Mathematical Theory of Quantitative Genetics. Oxford Univ. Press, Oxford, UK.

Campos, J. L., and B. Charlesworth. 2019. The effects on neutral variability of recurrent selective sweeps and background selection. Genetics 212: 287–303.

Campos, P. R. A., and L. M. Wahl. 2009. The effects of population bottlenecks on clonal interference, and the adaptation effective population size. Evolution 63: 950–958.

Campos, P. R. A., and L. M. Wahl. 2010. The adaptation rate of asexuals: deleterious mutations, clonal interference and population bottlenecks. Evolution 64: 1973–1983.

Charlesworth, B. 2013a. Background selection 20 years on. J. Hered. 104: 161–171.

Charlesworth, B. 2013b. Stabilizing selection, purifying selection, and mutational bias in finite populations. Genetics 194: 955–971.

Charlesworth, B. 2020. How long does it take to fix a favorable mutation, and why should we care? Amer. Nat. 195: 753–771.

Charlesworth, D., B. Charlesworth, and M. T. Morgan. 1995. The pattern of neutral molecular variation under the background selection model. Genetics 141: 1619–1632.

Charlesworth, B., and K. Jain. 2014. Purifying selection, drift, and reversible mutation with arbitrarily high mutation rates. Genetics 198: 1587–1602.

Cockerham, C. C. 1984. Drift and mutation with a finite number of allelic states. Proc. Natl. Acad. Sci. USA 81: 530–534.

Bulmer, M. 1991. The selection-mutation-drift theory of synonymous codon usage. Genetics 129: 897–907.

Desai, M. M., and D. S. Fisher. 2007. Beneficial mutation selection balance and the effect of linkage on positive selection. Genetics 176: 1759–1798.

Kingsolver, J. G., and S. E. Diamond. 2011. Phenotypic selection in natural populations: what limits directional selection? Am. Nat. 177: 346–357.

Eshel, I., and M. W. Feldman. 1970. On the evolutionary effect of recombination. Theor. Popul. Biol. 1: 88–100.

Gale, J. S. 1990. Theoretical Population Genetics. Unwin Hyman, London, UK.

Gerrish, P. J., and R. E. Lenski. 1998. The fate of competing beneficial mutations in an asexual population. Genetica 102/103: 127–144.

Gessler, D. D. 1995. The constraints of finite size in asexual populations and the rate of the ratchet. Genet. Res. 66: 241–253.

Good, B. H., A. M. Walczak, R. A. Neher, and M. M. Desai. 2014. Genetic diversity in the interference selection limit. PLoS Genet. 10: e1004222.

Goyal, S., D. J. Balick, E. R. Jerison, R. A. Neher, B. I. Shraiman, and M. M. Desai. 2012. Dynamic mutation-selection balance as an evolutionary attractor. Genetics 191: 1309–1319.

Jain, K. 2019. Interference effects of deleterious and beneficial mutations in large asexual populations. Genetics 211: 1357–1369.

Jain, K., and S. John. 2016. Deterministic evolution of an asexual population under the action of beneficial and deleterious mutations on additive fitness landscapes. Theor. Popul. Biol. 112: 117–125.

John, S., and K. Jain. 2015. Effect of drift, selection and recombination on the equilibrium frequency of deleterious mutations. J. Theor. Biol. 365: 238–246.

Johnson, T., and N. H. Barton. 2002. The effect of deleterious alleles on adaptation in asexual populations. Genetics 162: 395–411.

Kim, Y., and W. Stephan. 2000. Joint effects of genetic hitchhiking and background selection on neutral variation. Genetics 155: 1415–1427.

Kimura, M. 1969. The number of heterozygous nucleotide sites maintained in a finite population due to steady flux of mutations. Genetics 61: 893–903.

Kimura, M. 1983. The Neutral Theory of Molecular Evolution. Cambridge Univ. Press, Cambridge, UK.

Kimura, M, and J. F. Crow, 1964. The number of alleles that can be maintained in a finite population. Genetics 49: 725–738.

Kimura, M., T. Maruyama, and J. F. Crow. 1963. The mutation load in small populations. Genetics 48: 1303–1312.

Latter, B. D. 1970. Selection in finite populations with multiple alleles. II. Centripetal selection, mutation, and isoallelic variation. Genetics 66: 165–186.

Lande, R. 1975. The maintenance of genetic variability by mutation in a polygenic character with linked loci. Genetical Research 26: 221–235.

Li, W. H. 1987. Models of nearly neutral mutations with particular implications for non-random usage of synonymous codons. J. Mol. Evol. 24: 337–345.

Long, H., W. Sung, S. Kucukyildirim, E. Williams S., W. Guo, C. Patterson, C. Gregory, C. Strauss, C. Stone, C. Berne, D. Kysela, W. R. Shoemaker, M. Muscarella, H. Luo, J. T. Lennon, Y. V. Brun, and M. Lynch. 2017. Evolutionary determinants of genome-wide nucleotide composition. Nature Ecol. Evol. 2: 237–240.

Lynch, M. 2011. The lower bound to the evolution of mutation rates. Genome Biol. Evol. 3: 1107–1118.

Lynch, M. 2018. Phylogenetic diversification of cell biological features. eLife 7: e34820.

Lynch, M. 2020. The evolutionary scaling of cellular traits imposed by the drift barrier. Proc. Natl. Acad. Sci. USA 117: 10435–10444.

Lynch, M., M. Ackerman, J.-F. Gout, H. Long, W. Sung, W. K. Thomas, and P. L. Foster. 2016. Genetic drift, selection, and evolution of the mutation rate. Nature Rev. Genetics 17: 704–714.

Lynch, R. BÖrger, D. Butcher, and W. Gabriel. 1993. Mutational meltdowns in asexual populations. J. Heredity 84: 339–344

Lynch, M., and W. G. Hill. 1986. Phenotypic evolution by neutral mutation. Evolution 40: 915–935.

Lynch, M., and W.-C. Ho. 2020. The limits to estimating population-genetic parameters with temporal data. Genome Biol. Evol. 12: 443–455.

Lynch, M., and B. Trickovic. 2020. A theoretical framework for evolutionary cell biology. J. Mol. Biol. 432: 1861–1879.

Lynch, M., B. Trickovic, and C. P. Kempes. 2022. Evolutionary scaling of maximum growth rates with organism size. Scientific Reports (in press).

McVean, G., and B. Charlesworth. 1999. A population genetic model for the evolution of synonymous codon usage: patterns and predictions. Genet. Res. 74: 145–158.

Pénisson S., T. Singh, P. Sniegowski, and P. Gerrish. 2017. Dynamics and fate of beneficial mutations under lineage contamination by linked deleterious mutations. Genetics 205: 1305–1318.

Walsh, J. B., and M. Lynch. 2018. Evolution and Selection of Quantitative Traits. Oxford Univ. Press, Oxford, UK.

## Literature Cited

Gale, J. S. 1990. Theoretical Population Genetics. Unwin Hyman, London, UK.

Kimura, M. 1983. The Neutral Theory of Molecular Evolution. Cambridge Univ. Press, Cambridge, UK.

Rouzine, I. M., E. Brunet, and C. O. Wilke. 2008. The traveling wave approach to asexual evolution: Muller’s ratchet and speed of adaptation. Theor. Popul. Biol. 73: 24–46.

